# Chk2 homologue Mek1 limits Exo1-dependent DNA end resection during meiotic recombination in *S. cerevisiae*

**DOI:** 10.1101/2024.04.12.589255

**Authors:** Jennifer Grubb, Douglas K. Bishop

**Affiliations:** Department of Radiation and Cellular Oncology/Department of Molecular Genetics and Cell Biology, University of Chicago, 920 E 58^th^ Street, CLSC 817, Chicago, IL 60637

**Keywords:** Chk2, Exo1, Mek1, DNA-end resection, meiosis, recombination

## Abstract

The conserved Rad2/XPG family 5’—3’ exonuclease, Exonuclease 1 (Exo1), plays many roles in DNA metabolism including during resolution of DNA double strand breaks (DSBs) via homologous recombination. Prior studies provided evidence that the end-resection activity of Exo1 is downregulated in yeast and mammals by Cdk1/2 family cyclin-dependent and checkpoint kinases, including budding yeast kinase Rad53 which functions in mitotic cells. Here we provide evidence that the master meiotic kinase Mek1, a paralogue of Rad53, limits 5’-3’ single strand resection at the sites of programmed meiotic DNA breaks. Mutational analysis suggests that the mechanism of Exo1 suppression by Mek1 differs from that of Rad53.

**Article Summary:** Meiotic recombination involves formation of programmed DNA double strand breaks followed by 5’ to 3’ single strand specific resection by nucleases including Exo1. We find that the activity of budding yeast Exo1 is downregulated during meiotic recombination by the master meiotic kinase Mek1. The mechanism of downregulation of Exo1 by Mek1 in meiosis does not depend on the same phospho-sites as those used by the mitotic kinase Rad53, a relative of Mek1 that downregulates Exo1 in mitosis.

## Introduction

Exonuclease 1 (Exo1) is a 5’-3’ strand specific double strand DNA exonuclease and a member of the Rad2/XPG family of eukaryotic exonucleases (Szankasi and Smith 1992; Fiorentini *et al*. 1997). Exo1 plays roles in many DNA metabolic pathways including nucleotide mismatch repair, nucleotide and base excision repair, DNA replication stress tolerance, telomere maintenance, DNA damage checkpoint signaling, and micro-homology mediated DNA end-joining (Symington 2016; Yan *et al*. 2021). In addition, Exo1 plays critical roles in homologous recombination during mitosis and meiosis.

Exo1 plays an enzymatic role at sites of DNA double strand breaks (DSBs) to form the ssDNA tracts onto which the recombination machinery assembles (Zakharyevich *et al*. 2010; Mimitou *et al*. 2017). Exo1 also plays a non-enzymatic role during budding yeast meiosis that is required for the resolution of Holliday junction intermediates into reciprocal crossover recombinants (Zakharyevich *et al*. 2010; Hunter 2011; Sanchez *et al*. 2020; Gioia *et al*. 2023).

Exo1’s activity is highly regulated by various post-translational modifications including acetylation, ubiquitination, SUMOylation, and phosphorylation (Symington 2016; Sertic *et al*. 2020). Phosphorylation of Exo1 is controlled by DNA damage checkpoint kinases including Mec1 and Tel1 in budding yeast (Carballo *et al*. 2008; Morin *et al*. 2008a; Bolderson *et al*. 2010; Tomimatsu *et al*. 2014; Morafraile *et al*. 2020a). Checkpoint-dependent phosphorylation of Exo1 has been reported to occur via pathways in which a phosphatidylinositol 3-kinase-related kinase (PIKK) directly phosphorylates Exo1 as well as via pathways involving indirect PIKK-mediated activation of effector kinases of the Cdk1,2 family, which also downregulate Exo1’s activity during the G1 stage of the cell cycle.

A previous study in budding yeast identified four sites on Exo1 that are phosphorylated by the Chk2 orthologue Rad53 (Morin *et al*. 2008a). Mutation of the four Rad53 phosphorylation sites on Exo1 altered cell growth and viability in a manner consistent with the notion that the relevant phosphorylated forms of Exo1 are less active than the corresponding non-phosphorylated forms. In another study, mutation of a set of 23 Rad53 phosphorylation sites provided evidence that 911 checkpoint signaling via Rad53 limits Exo1 activity to protect cells from the impact of replication stress (Morafraile *et al*. 2020a).

The budding yeast kinase Mek1 is a meiosis specific paralogue of Rad53 (Hollingsworth and Gaglione 2019). Once activated, Mek1 regulates several meiotic processes. First, Mek1 blocks the homology search and strand exchange activity of the mitotic homologous recombinase Rad51 by two mechanisms including phosphorylation-dependent stabilization of the Rad51 inhibitor Hed1 (Tsubouchi and Roeder 2006; Callender *et al*. 2016) and by inhibition of the Rad51 accessory protein Rad54 (Niu *et al*. 2009). This downregulation of Rad51’s activity converts Rad51 to an accessory factor for the meiosis-specific recombinase Dmc1 (Cloud *et al*. 2012). Mek1 also phosphorylates the synaptonemal complex protein Zip1 (Chen *et al*. 2015). Another notable activity of Mek1 is its role in the meiotic recombination checkpoint which arrests meiotic progression until recombination mediated repair of programmed meiotic DNA breaks is complete (Lydall *et al*. 1996; Xu *et al*. 1997). When activated by the upstream checkpoint kinase Mec1, Mek1 promotes meiotic prophase arrest by inhibitory phosphorylation of Ndt80, a master transcription factor required for meiotic progression from prophase to metaphase I (Chen *et al*. 2018). Mek1 is also required for a regulatory process that promotes recombination between homologous chromatids, as opposed to sister chromatids (Niu *et al*. 2005, 2007; Kim *et al*. 2010).

Here we show that Mek1 limits end-resection at the sites of programmed meiotic DNA breaks in budding yeast. The hyper-resection seen in Mek1 deficient cells depends on Exo1, and Mek1 promotes phosphorylation of Exo1. The results provide evidence for a mechanism triggered by single strand DNA-dependent checkpoint signaling that restricts the length of meiotic resection tracks.

## Results and Discussion

### The staining intensity of RPA foci increases when Mek1 activity is suppressed

To determine the role of Mek1 in DNA-end resection, we used a strain that expresses a “druggable” mutant form of the kinase Mek1*, mek1-as1*, which retains its function under standard growth conditions, but is inhibited by the ATP analogue 1-Naphthl-PP1 (1-NA-PP1) (Bishop *et al*. 2001) (Wan *et al*. 2004). We examined recruitment of RPA to sites of meiotic breaks by immunostaining of surface spread nuclei prepared at various times after induction of meiosis. Using this system, we found that inhibition of Mek1-as1 with 1-NA-PP1 enhanced overall RPA staining 1.8-fold (Figure 1). Interestingly, while the average staining intensity of foci was increased about 3-fold, the average number of foci per nucleus was reduced 2-fold. We showed previously that a *rad51Δ* mutant displays a similar RPA staining pattern to that seen here for *mek1-as1* (Gasior *et al*. 1998). Given that *rad51Δ* mutants display hyper-resection (Shinohara *et al*. 1992), the similarities in RPA staining suggested that inhibition of Mek1 might also cause hyper-resection.

**Figure 1.**
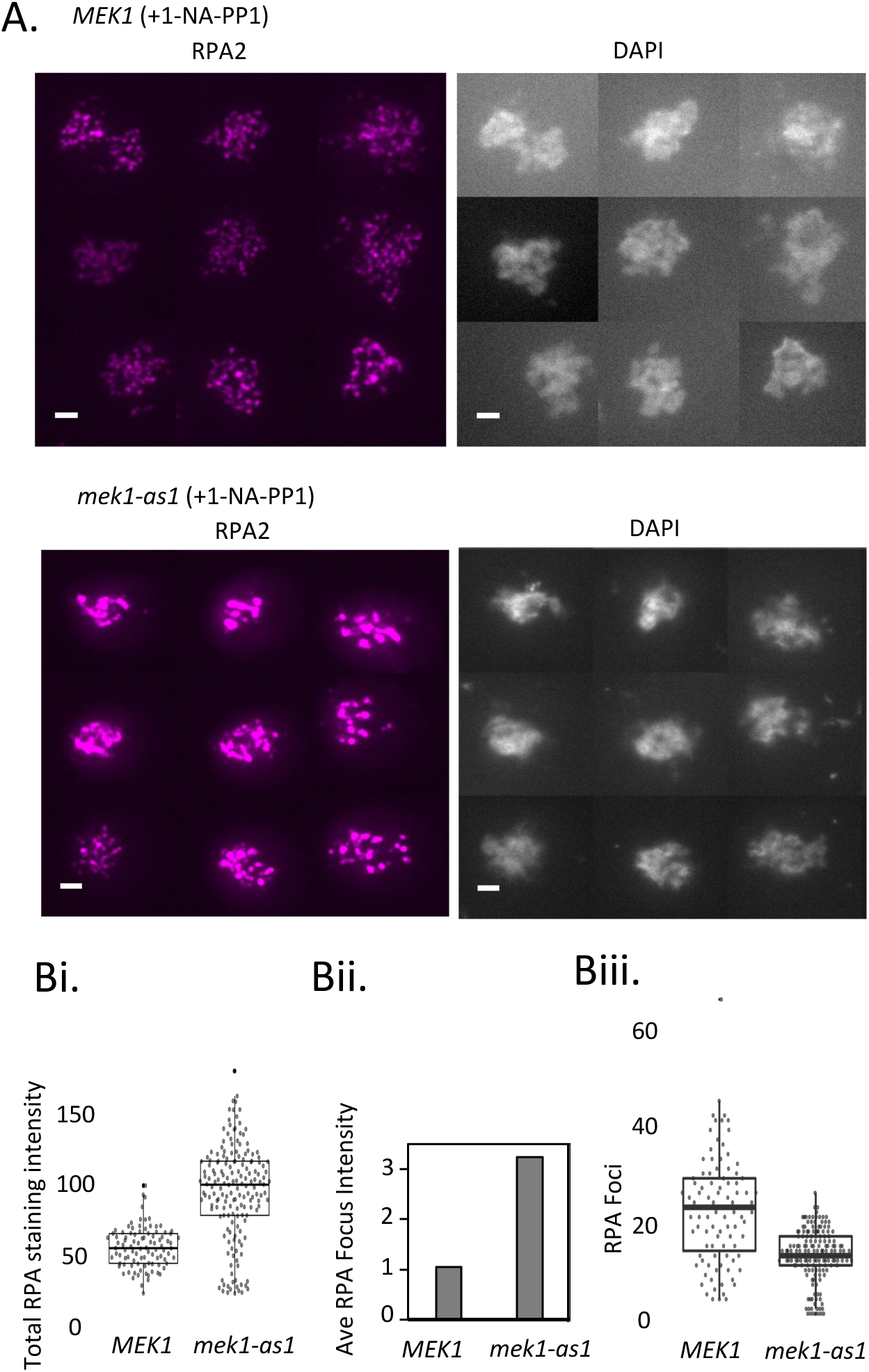
RPA focus staining intensity increases when Mek1 activity is suppressed. Immunostaining of spread meiotic nuclei. Spread nuclei of *MEK1^+,^ rad51-II3A, dmc1-II2A, ndt80* (DKB6219) and *mek1-as1, rad51-II3A, dmc1-II2A, ndt80* (DKB6220) were prepared following 9 hours incubation in SPM and indirectly immunostained for RPA2. 1 µM 1-NA-PP1 was added to all cultures at time=0 hrs**. A.** Representative examples of spread nuclei. Size bar=2 microns. **B.** i. Quantification of total staining intensity of unselected spread nuclei. The box represents the upper and lower quartiles, and the central horizontal line the median. **Bi.** Quantification of total staining intensity of unselected spread nuclei. The box represents the upper and lower quartiles, and the central horizontal line the median. **Bii.** Quantification of the average staining intensity of foci. **Biii.** Quantification of the average number of foci per nucleus.

### Mek1 suppresses hyper-resection at the *HIS4::LEU2* recombination hotspot

To determine directly if Mek1 controls the amount of DNA end-resection during meiotic recombination, we made use of the well-characterized meiotic hotspot *HIS4::LEU2* (Cao *et al*. 1990; Lao *et al*. 2013). To facilitate detection of changes in resection activity, we used a strain containing mutations that allow formation, but not repair, of resected DNA ends. The strain carries alleles that express catalytically dead versions of recombinases Dmc1 and Rad51 (Cloud *et al*. 2012), as well as a *ndt80* null allele which prevents progression out of meiotic prophase (Xu *et al*. 1995). The strain also carries the *mek1-as1* allele to allow inhibition of Mek1 activity. Hereafter, we refer to this *mek1-as1, dmc1-II2A, rad51-II3A, ndt80* strain background as the “DSB-stuck” background.

We examined the structure of DNA breaks at the recombination hotspot *HIS4::LEU2* (Cao *et al*. 1990). We found that while resection of the DNA breaks formed at *HIS4::LEU2* was readily detectable as a signal composed of indirectly end-labelled species of variable mobility, inhibition of Mek1 activity with 1-NA-PP1 in the same DSB-stuck strain resulted in dramatically pronounced hyper-resection; the average mobity of DSB fragments detected in cells lacking Mek1 activity was consistently greater than that of fragments generated by cells with Mek1 activity (Figure 2). Thus, Mek1 functions to down regulate the 5’ to 3’ ssDNA resection. Because the DSB fragments observed in Mek1-deficient cells were more heterogenous, it was unclear if their total number was similar to that in cells with Mek1 activity. To confirm that DSB formation was not reduced by Mek1 inhibition, we used a *rad50S* mutation which blocks break resection. As reported previously (Xu *et al*. 1997), we saw no reduction in DSB formation in Mek1-deficient *rad50S* cells indicating that the relatively faint DSB signal observed in Mek1 deficient cells was the result of variable degrees of hyper-resection rather than a reduction in the efficiency of DSB formation (Figure S1).

**Figure 2.**
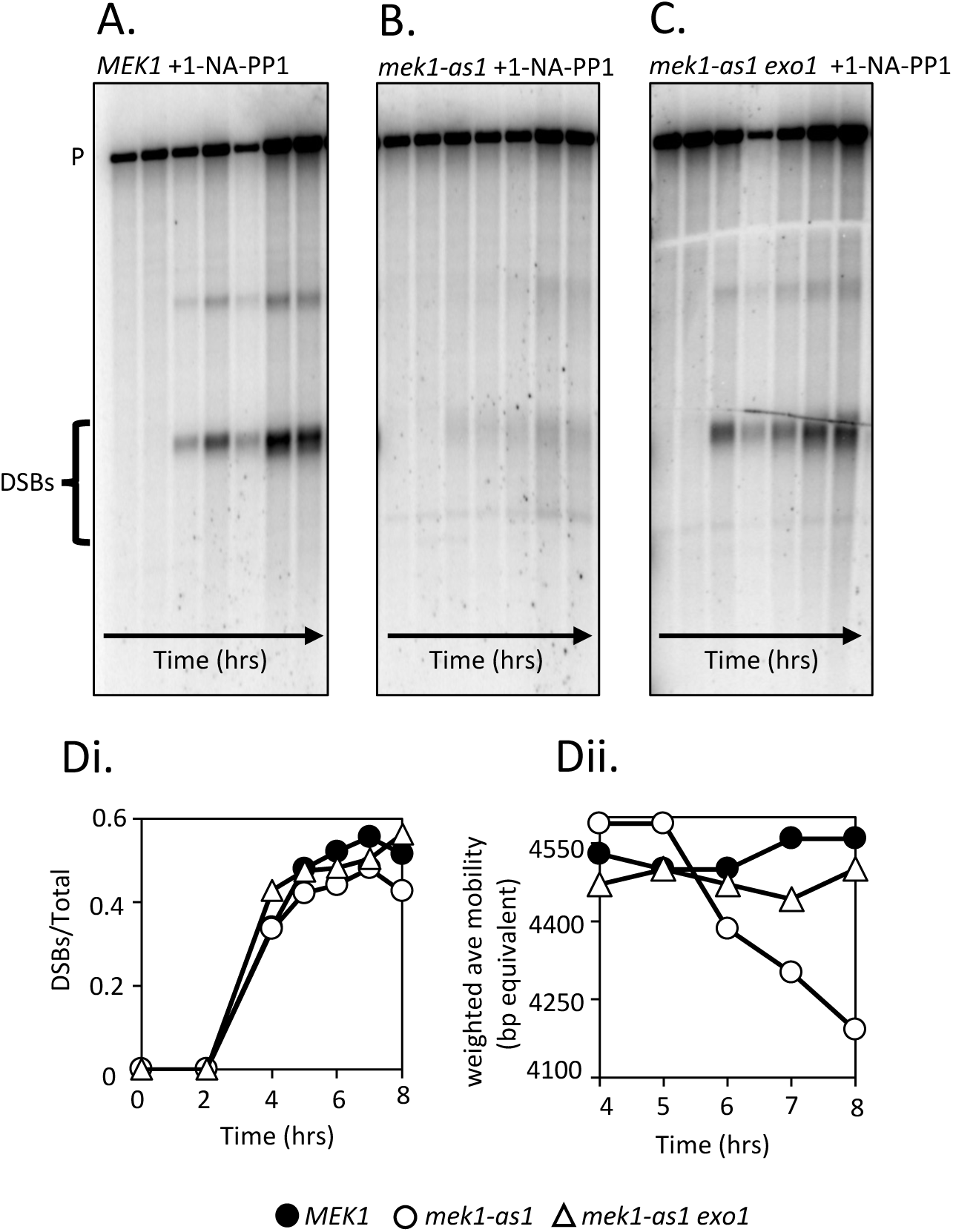
Mek1 limits DNA end resection. Southern analysis of DSB formation at the *HIS4::LEU2* hotspot. P=parental PstI fragment. DSBs=break fragments. Time points=0,2,4,5,6,7,8, and 9 hours. **A.** *MEK1^+,^ rad51-II3A, dmc1-II3A, ndt80* (DKB6219). **B***. mek1-as1, rad51-II3A, dmc1-II3A, ndt80* (DKB6220). C. *mek1-as1, exo1Δ, rad51-II3A, dmc1-II3A, ndt80* (DKB6337). 1 µM 1-NA-PP1 added to all cultures at time=0 hrs. . **Di.** Quantification of DSB signal/total DNA **Dii.** (**Note:** Fragments are expected to be resected 5’-3’ with ssDNA strands ending 3’ at the site of the initial breaking remaining full length. Thus, lengths of resection tracts are predicted to be greater than the difference in size of fully duplex markers. Nonetheless, we opted to use the mobility of fully duplex DNA as a metric for the degree of resection to allow comparison with results of other studies of resection of the same site.)

### Exo1 is required for the hyper-resection observed in Mek1-defective cells

Previous studies showed Exo1 is primarily responsible for long-range 5’-3’ ssDNA resection at meiotic DSB sites (Manfrini *et al*. 2010; Zakharyevich *et al*. 2010). We therefore asked if Exo1 activity was responsible for the hyper-resection we observed in Mek1-defective DSB stuck cells. We found that deletion of the *EXO1* gene in the DSB stuck background eliminated the hyper-resection observed upon Mek1 repression (Figure 2). This result provides evidence that Mek1-dependent repression of hyper-resection in DSB stuck cells results from repression of Exo1. One notable feature of the DSB signal observed in Mek1-deficient *exo1* cells was a striking similarity to that from Mek1-proficient *EXO1^+^* cells. Previous work in vegetative cells has shown that the Dna2 nuclease functions with the Sgs1 helicase to resect DNA ends in a pathway that is largely redundant with Exo1. These earlier findings raised the possibility that Dna2-Sgs1 is responsible for residual resection in *mek1 exo1* defective cells (Symington 2016; Gnügge and Symington 2021). Consistent with this, a prior study in *dmc1* cells provided evidence that Sgs1 can contribute significant resection activity in the absence of Exo1 (Manfrini *et al*. 2010). On the other hand, a second study using *DMC1^+^* cells did not detect resection activity for Sgs1 even in an *exo1* mutant background (Zakharyevich *et al*. 2010). Thus, additional studies will be required to determine which nuclease is active in *mek1 exo1* double mutants in our DSB stuck background.

### Meiotic recombination checkpoint signaling is required for Mek-1 dependent repression of Exo1

Previous work established that Mek1 kinase activity is activated by the checkpoint kinase Mec1, a homolog of the mammalian “911” damage-sensitive kinase ATR. Mec1 promotes checkpoint control during meiotic recombination (Lydall *et al*. 1996). Mec1 is recruited to tracts of ssDNA where it functions in conjunction with the so-called “911 clamp” complex to induce phosphorylation of downstream effector proteins. In budding yeast, the clamp complex is composed of 3 proteins; Ddc1, Mec3, and Rad17 (Roeder and Bailis 2000). The activity of Mec1 depends on all the clamp proteins. To determine if activation of Mek1 by 911 signaling is required to repress Exo1 dependent resection, we examined the impact of a *rad17* mutation in the DSB stuck background. We found that a *rad17* mutant displayed hyper-resection even in the presence of functional Mek1, consistent with the known requirement for Rad17 in Mek1 activation (Figure 3). Interestingly, the degree of hyper-resection was more pronounced when Mek1 activity was repressed in *rad17* cells compared *RAD17^+^* cells. This result suggests Rad17 has both Mek1-dependent and Mek1-independent functions that limit resection.

**Figure 3.**
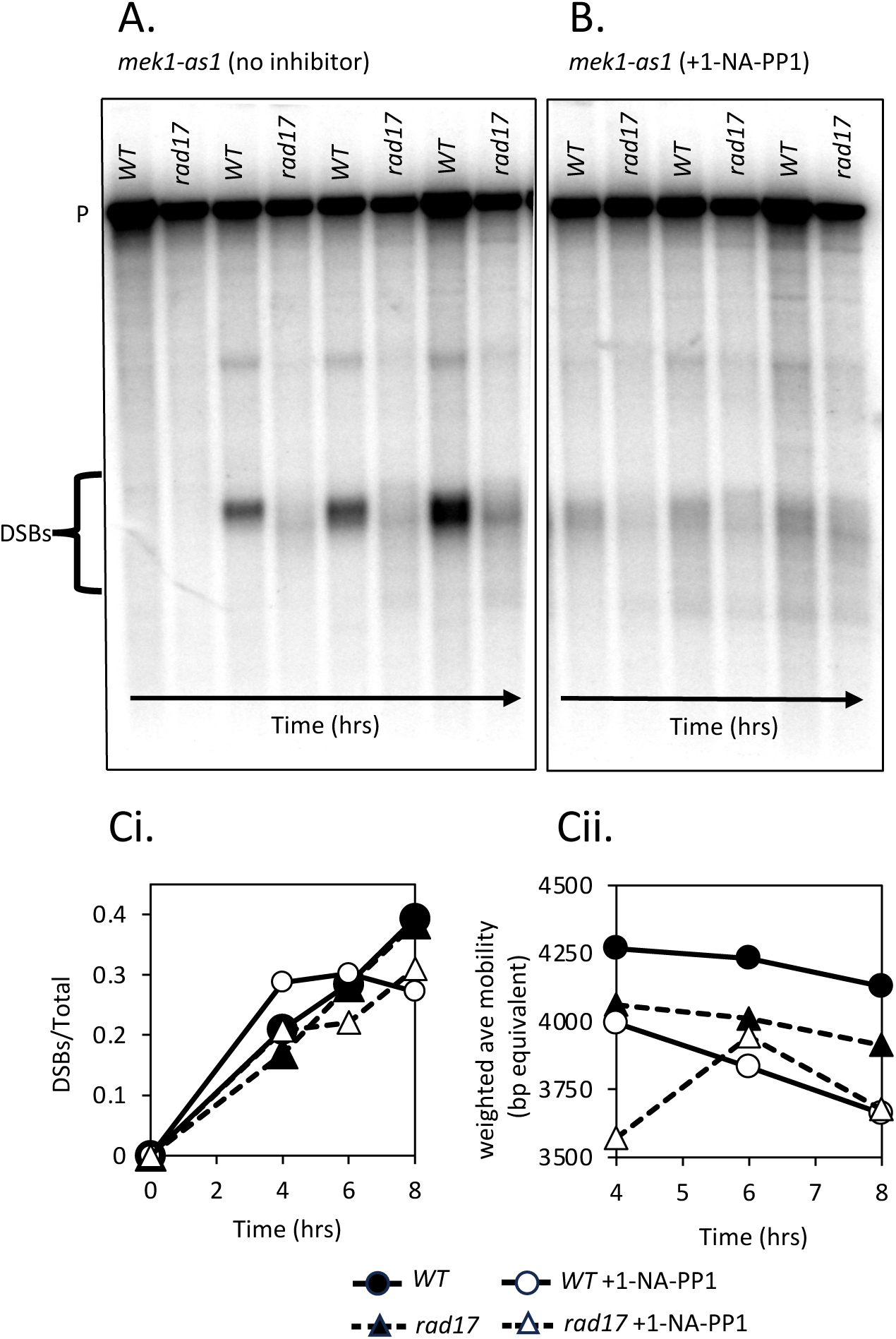
Rad17 limits DNA end resection. Southern analysis of DSB formation at the *HIS4::LEU2* hotspot. P=parental PstI fragment. DSBs=break fragments. Time points=0,4,6,and 8 hours. **A.** DMSO only (no inhibitor) **B.** 1µM 1-NA-PP1 was added at time=0hrs. **Ci.** Quantification of DSB signal/total DNA **Cii.** Weighted average of DSB fragment lengths based on mobility of fully duplex DNA size markers. See note in the legend of Figure 2. Strains used: *mek1-as1, rad51-II3A, dmc1-II3A, ndt80, EXO1-13XcMyc* (DKB6800) and *mek1-as1, rad51-II3A, dmc1-II3A, rad17Δ, EXO1-13XcMyc* (DKB6903).

### Exo1 is phosphorylated by Mek1

We next used Western blot analysis to determine if Exo1 showed evidence of phosphorylation by Mek1. Whole cell extracts were prepared from meiotic cultures 4hrs after transfer of cells to meiosis-inducing sporulation medium and analyzed using a gel system that enhances the mobility difference between phosphorylated and unphosphorylated protein forms (see Methods and Materials). In the strains used, Exo1 was tagged with a 13XcMyc epitope and detected using anti-cMyc antibodies (Longtine *et al*. 1998). This analysis revealed that Exo1 migrates as a closely spaced doublet during meiosis in DSB-stuck cells that express Mek1 activity (Figure 4). Treating extracts with lambda phosphatase prior to electrophoretic separation resulted in a dramatic reduction in the intensity of the slower migrating band of the Exo1 doublet, consistent with that band containing a phosphorylated form of the protein. Importantly, the intensity of the slower migrating band was also dramatically reduced when Mek1 activity was inhibited. These results indicate that Mek1 kinase is required for the normal level of phosphorylation of Exo1.

**Figure 4.**
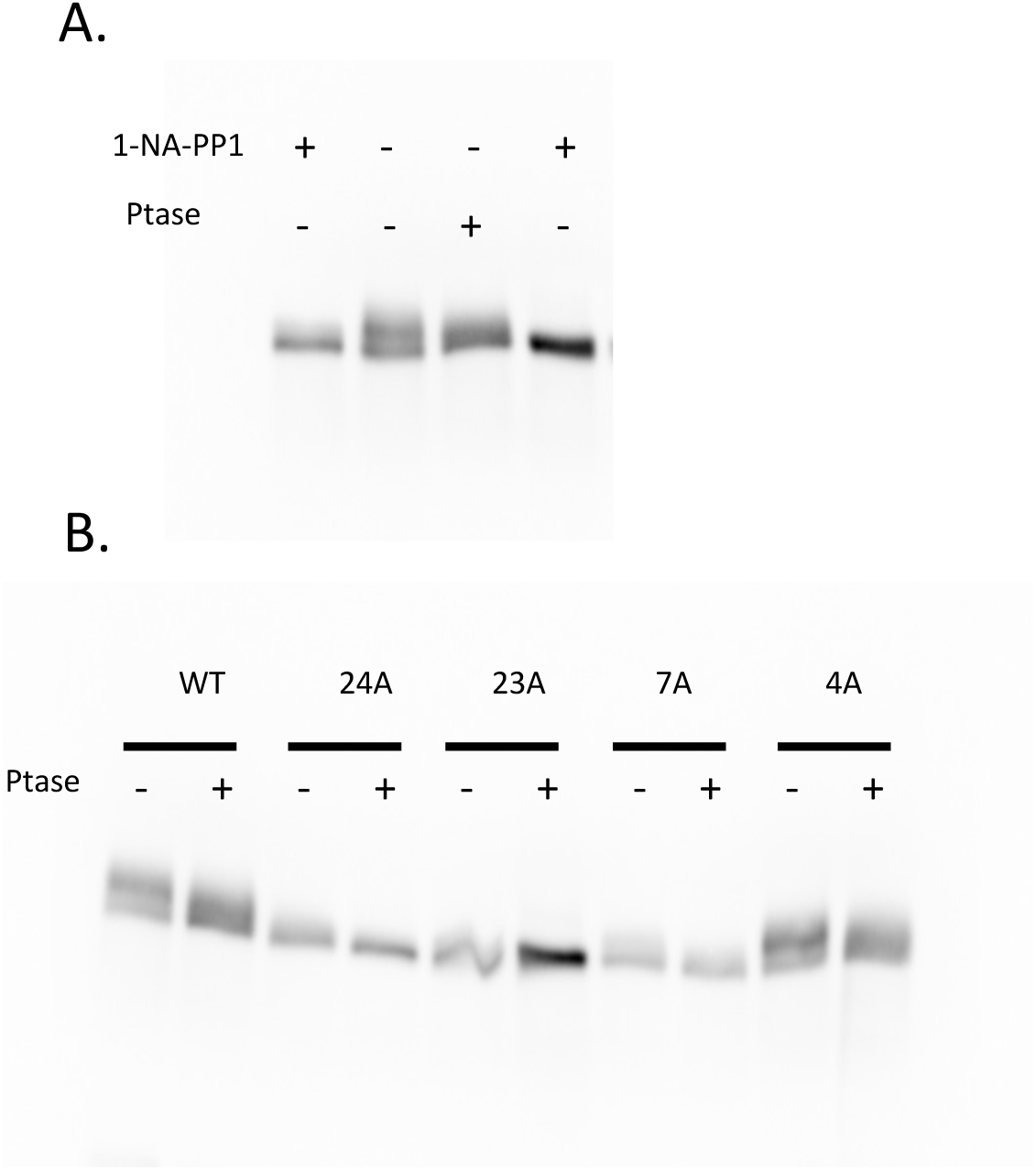
Exo1 displays Mek1-dependent phosphorylation. **A.** Western blot analysis of Exo1. Lysates were prepared from a 4hr meiotic timepoint of a DKB6800 culture. For lanes marked -, no inhibitor was added. For lanes marked +, 1 µM 1-NA-PP1 added at time=0hrs. TCA ppts were incubated with or without lambda phosphatase as indicated. **B.** Western blot analysis of Exo1 phospho site mutants. Strains used were *EXO1-13XcMyc* =DKB6800, *exo1-24A-13XcMyc*=DKB6810, *exo1-23A-13XcMyc* =DKB6764, *exo1-7A-13XcMyc* =DKB6613, *exo1-4S::A-13XcMyc*=DKB6611. Blots were probed with anti-cMyc antibodies.

### Analysis of Exo1 phosphorylation sites by mass spectrometry

To identify sites in Exo1 that are subject to Mek1 dependent phosphorylation, we analyzed Exo1 peptides by mass spectrometry (see Methods and Materials). Exo1 protein was purified from both *MEK1^+^* and *mek1Δ* meiotic cells. Three phospho-sites were detected in *MEK1^+^* but not *mek1Δ* cells (S692, S587, and S663). An additional phospho-site, S431, was shared by Exo1 purified from both *MEK1^+^* and *mek1Δ* cells.

### Mutational analysis of Exo1 phosphorylation sites

We next carried out experiments to identify additional phospho-sites in Exo1 that are required for its repression. All mutants examined are described in detail in Table S1. We took advantage of 3 prior studies in designing mutations at potential phosphorylation sites. First, a previous study on Exo1 phosphorylation in mitotic cells showed that the mitotic kinase Rad53, a paralogue of Mek1, phosphorylates Exo1 at 4 sites: S372, S567, S587, and S692 (Morin *et al*. 2008b). We therefore examined an Exo1 mutant in which all codons corresponding to these sites was changed to code for alanine (*exo1-4S::A*).

Another prior study had showed Mek1 prefers to phosphorylate threonine embedded in the consensus sequence RXXT (Suhandynata *et al*. 2016). Exo1 contains three such consensus sequences at T184, T249, and T558. We therefore constructed an allele of Exo1 carrying 7 mutated sites, the 4 phosphorylated in mitosis, and the 3 Mek1 consensus sites (*exo1-7A*). Two of the phosphorylation sites we detected by mass spectrometry were not represented among these 7 mutated sites in *exo1-7A*, therefore we created *exo1-8A* (adding S663) and *exo1-9A* (adding S663 and S431). Third, an earlier study of Exo1 phosphorylation made use of an allele in which 23 sites of potential phosphorylation were mutated (*exo1-23A*) (Morafraile *et al*. 2020b). One of the phosphorylation sites we detected by mass spectrometry (S663) was not represented among the 23 mutated in *exo1-23A* and this site was mutated to alanine to generate *exo1-24A* (Figure 4, Table S1). Additional relevant sites that are not represented in *exo1-24A* include 2 Mek1 consensus sites (T184, T558) and 2 sites phosphorylated in mitosis (S372, S567). We therefore created an additional Exo1 mutant that contained those 4 missing sites (*exo1-28A*). Finally, we added one additional mutation at a site we found to be phosphorylated in both *MEK1^+^* and *mek1Δ* cells (*exo1-29A*).

A subset of mutant strains containing different combinations of potential phosphorylation site mutations were compared to a strain expressing Exo1-WT using the Western assay. Exo1-4S::A strain showed a similar level of the phosphor-specific Exo1 band to that seen in Exo1-WT cells indicating that the sites of mitotic phosphorylation by Rad53 are largely dispensable for the phosphorylation-dependent mobility shift seen in Mek1 expressing cells. In contrast, the *exo1-7A*, *exo1-23A*, and *exo1-24A* mutants all showed a dramatic reduction in the intensity of the phospho-specific band. Given that two of the mek1 consensus sites were not mutated in exo1-23 and exo1-24, the results together imply that phosphorylation at T249 is responsible for most of the observed mobility shift (Figure 4).

We predicted that blocking Mek1’s ability to phosphorylate Exo1 by mutating the relevant phosphor-sites would result in hyper-resection in Mek1 expressing cells. However, for each of the *exo1* mutants examined, the appearance of the DSB fragments at *HIS4::LEU2* was either unchanged or the amount of resection was reduced, presumedly as a consequence of defects conferred by the mutations that are unrelated to phosphosite function (Figure 5, Figure S2). These results are consistent with at least two possibilities. First, Mek1 may be capable of inhibiting Exo1 activity by phosphorylating sites that were not mutated in *exo1-29A*. Alternatively, Mek1 may repress Exo1 activity by a mechanism that is independent of its ability to directly phosphorylate the nuclease.

**Figure 5.**
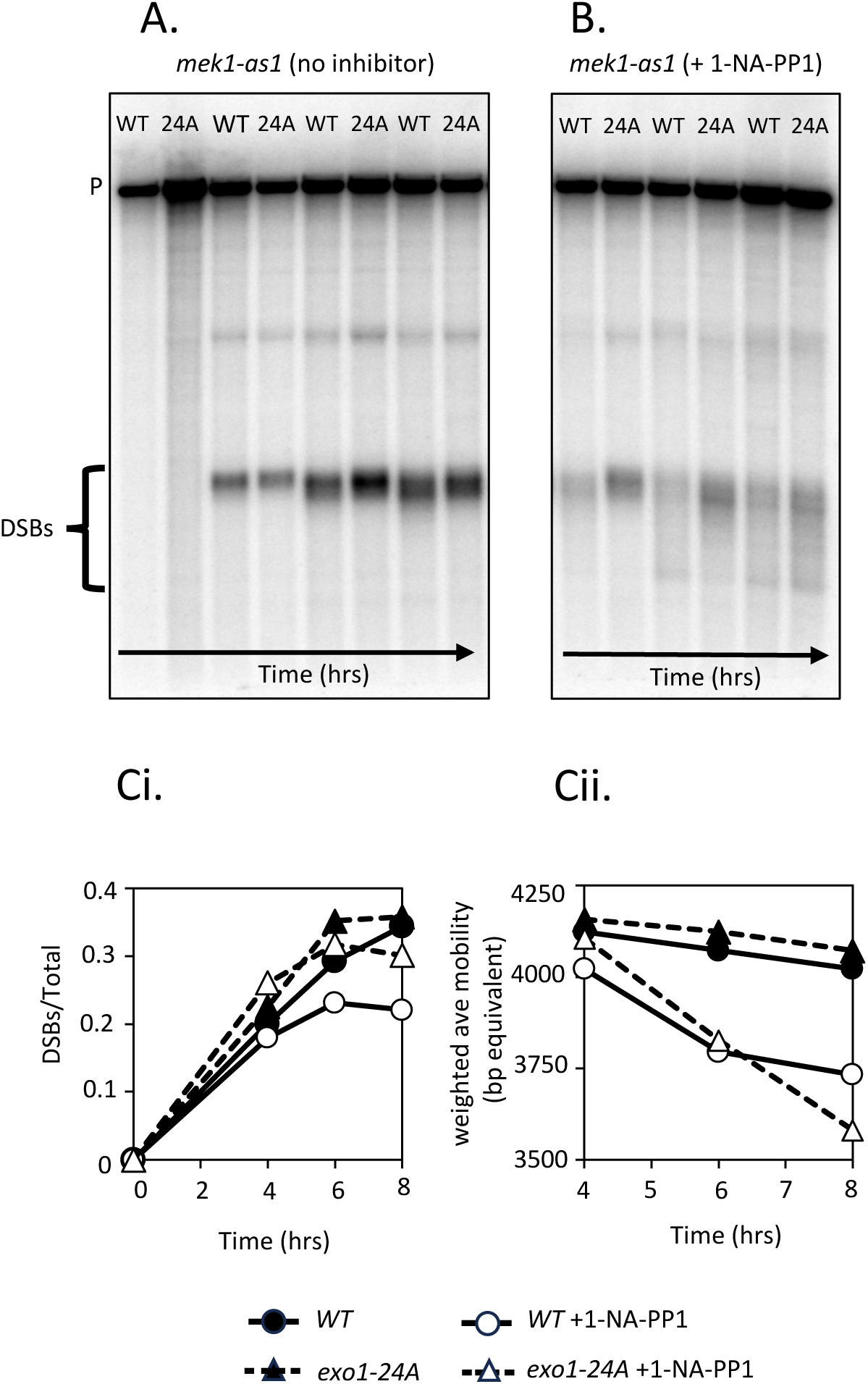
Mutant analysis of Exo1 phosphosites. Southern analysis of DSB formation at the *HIS4::LEU2* hotspot. P=parental PstI fragment. DSBs=break fragments. Time points=0,4,6 and 8 hours. Strains used were *mek1-as1, rad51-II3A, dmc1-II3A, ndt80, EXO1-13XcMyc* (DKB6800) and *mek1-as1, rad51-II3A, dmc1-II3A, ndt80, exo1-24A-13XcMyc* (DKB6810). Note that the *exo1-24A-13XcMyc* strain displays modestly reduced resection both with and without Mek1 inhibition. **A.** DMSO only (no inhibitor) was added at time=0hrs. **B.** 1µM 1-NA-PP1 was added at time=0hrs. **Ci.** Quantification of DSB signal/total DNA **Cii.** Weighted average of DSB fragment lengths based on mobility of fully duplex DNA size markers. See note in the legend of Figure 2.

### Possible functions of downregulation of Exo1 during meiosis

Demonstration that the activity of Exo1 is actively downregulated during meiosis raises the question of whether the range of resection tract lengths that occur during unperturbed meiosis is optimized to promote favorable recombination outcomes. A previous study showed that relatively long tracts of ssDNA can engage multiple recombination donors simultaneously leading to genome rearrangement (Piazza *et al*. 2017). Other studies found that mutants which alter break repair can cause multi-chromatid joints to accumulate to higher-than-normal levels (Oh *et al*. 2008; Reitz *et al*. 2019). Given these observations, it is interesting to speculate that negative regulation of Exo1 activity may serve to limit recombination events involving more than 2 chromatids. It is also possible that Mek1’s role in promoting interhomolog, as opposed to intersister, recombination is functionally related to its role in limiting end-resection (Niu *et al*. 2005, 2009; Kim *et al*. 2010). Further studies will be required to determine the impact of altering resection tract lengths in recombination proficient cells without perturbing the many other regulatory roles of Mek1. It is also worth noting that regulation of Exo1’s nuclease activity by Mek1 raises the possibility that Mek1 may also regulate Exo1’s nuclease-independent function in promoting meiotic crossover recombination.

In conclusion, our results show that Mek1 downregulates Exo1 activity during meiosis and appears to do so by a mechanism that differs significantly from the previously described downregulation of Exo1 by Mek1’s mitotic paralogue Rad53.

## Materials and Methods

### Yeast strains

Strains used were isogenic to SK-1 and constructed by genetic crosses or by transformation. Genotypes and plasmids are listed in Supplementary Table 2 and 3. DNA constructs used for targeting mutations were made using the Gibson method (NEB, #E2621L) with details provided in the supplemental Materials and Methods.

### Induction of meiosis

Yeast cultures were induced to undergo meiosis as previously described with the following change: yeast was shifted into sporulation medium (SPM) when the OD_600_ was between 0.7-1 (Bishop 1994). Details of these methods are provided in the supplemental Materials and Methods. Mek1-as1 was inhibited with 1µM 1-NA-PP1 (Cayman Chemical, catalogue # 10954) in DMSO added to SPM medium immediately before incubation (0hr) (Wan *et al*. 2004).

### Immunostaining of spread nuclei and microscopy

Surface spreading and immunostaining of yeast chromosomes were performed as previously described (Grubb *et al*. 2015). Details regarding antibodies and imaging are provided in the supplemental Materials and Methods.

### Southern Blot analysis of DNA breaks

Southern blotting analysis of DSB fragments from the *HIS4::LEU2* hotspot was carried out using methods described previously (Lao *et al*. 2013).

### Lysates/Westerns

Total protein lysates were prepared by TCA precipitation using a previously described method (Knop *et al*. 1999) with alterations described in the supplemental Materials and Methods.

### Phosphatase treatment

TCA pellets were resuspended in PMP buffer (50mM HEPES, 100mM NaCl, 2mM DTT, 0.01% Brij 35, 1mM MnCl_2_) with 800 units Lambda phosphatase (New England Biolabs P0753S) in a final volume of 70μl and incubated at 30°C for 45min. 70μl of Buffer H (200mM Tris-HCl [pH 6.5], 8M urea, 5% SDS, 1mM EDTA, 0.02% bromophenol blue, and 5% - mercaptoethanol) was added to the mixture and the protein samples were denatured for 10min at 65°C, spun for 3min at full speed and the supernatant then subjected to Western analysis.

### Exo1 purification and mass spectrometry

Exo1 was purified from about 1g cells from 200ml cultures incubated for 4hrs under the previously described meiotic conditions. Protein was prepared from both *MEK1^+^* (DKB6713) and *mek1Δ* (DKB6704) strains. Details of the methods used for generating cleared lysates, immunoprecipitation, and gel purification are provided in the Supplemental Materials and Methods. In-gel digestion and analysis by LC-MS/MS was carried out at PTM Biolab LLC (Shevchenko *et al*. 2006). A description of the methods used by the company is quoted in the supplemental Materials and Methods.

## Data Availability Statement

All of the data used to confirm the results of this study are located within the manuscript, figures and tables. Yeast strains and plasmids used in this study are available upon request.

## Acknowledgments

We thank Gönen Memisoglu for helpful advice and reagents used for CRISPR-mediated mutation of *RAD17*. Thanks to Akira Shinohara for anti-rabbit RFA2 antibodies, Mónica Segurado for the *exo1-23A* allele, and Lorraine Symington for the *exo1-4S::A* allele. We also thank Lorraine Symington, Nancy Hollingsworth, and Neil Hunter for helpful advice during these studies. Special thanks to Yuen-Ling Chan and Brian Budke for technical help. Mass spectrometry analysis was carried out at PTM BIO LLC.

## Supplemental Methods and Materials

### Plasmid and strain construction

Proper targeting and allele verification were confirmed by PCR of genomic DNA followed by sequencing. All strains were generated by transformation or crosses in the SK-1 strain background.

The initial ***exo1Δ*** strain (DKB6328) was generated from strain DKB5420 using a targeting fragment from plasmid pNRB718. pNRB718 was created using the Gibson assembly method with the help of a kit (NEB, E2621L). The plasmid contains a 490bp homology domain directly upstream of Exo1 from SK-1 (394525-395014), a hygromycin B cassette (pRS41H) (Taxis and Knop 2006) and 490bp homology domain from SK-1 directly downstream of Exo1 (391925-392415). The targeting fragment was generated by endonucleolytic digestion with EcoRV and BamHI and transformed into yeast as previously described (Gietz and Schiestl 2007). Transformants were screened for Hygromycin resistance and checked for proper insertion via PCR.

To construct the ***exo1-4S::A*** mutant (DKB6589), a plasmid designated pNRB721 was constructed by introducing a 4.6kb fragment obtained by PCR amplification using primers from 218bp upstream to 214bp downstream of Exo1 using genomic DNA from *exo1-4S::A* containing yeast strain LSY3958-2C, a gift from Dr. Symington, as template (DKB6364). This fragment was cloned into vector pBluescript using the Gibson assembly method. Then the *HIS3* marker was swapped out via Gibson for a kanMX4 cassette (from pRS41K) (Taxis and Knop 2006). The targeting fragment from the resulting plasmid (pNRB743) was generated by endonucleolytic digestion with EcoRV and Xho1 and transformed into *exo1Δ* strains (DKB6559, DKB6564) as previously described (Gietz and Schiestl 2007). A parallel construction was carried out with a corresponding fragment of *EXO1^+^* to serve as control (pNRB744).

To construct the ***exo1-7A*** mutant (DKB6586), pNRB743 (*exo1-4S::A*) was used as vector and primers for Gibson assembly designed that included the 3 Mek1 consensus site changes (RXXT to RXXA) (Suhandynata *et al*. 2016). The specific residues mutated are T184A, T249A, and T558A. The resulting plasmid (pNRB749) was cut and transformed into *exo1Δ* strains as described above.

The ***exo1-23A*** mutant (DKB6759, pNRB800), was constructed by the same method as used for *exo1-4S::A* except the *exo1* fragment (4.8kb) was obtained by PCR using DNA from strain *exo1-23A,* NAT52, a gift from Dr. Segurado (Morafraile *et al*. 2020).

For the ***exo1-24A*** mutant (DKB6798, DKB6799, pNRB803, pNRB811), primers were used to generate a S663A mutation in plasmid pNRB800 (which carried *exo1-23A*) by the same methods described above.

To construct the ***exo1-8A*, *exo1-9A*, *exo1-28A***, and ***exo1-29A*** mutants (pNRB935, pNRB936, pNRB938, pNRB939), gene blocks supplied by IDT were created with the appropriate changes in the coding region of *EXO1* and a Gibson reaction was used to replace the WT *EXO1* sequence from pNRB807 and create a CEN ARS plasmid for each mutant. These plasmids were then transformed into DKB6775 (DKB7052, DKB7053, DKB7055, and DKB7056 respectively).

The ***rad17Δ*** allele was generated without a selectable marker by CRISPR-Cas9 mediated targeting as described previously (Anand *et al*. 2017). A guide RNA sequence (CTGATATCTAAATCTCAGCT) was cloned into plasmid (bRA89) (Anand *et al*. 2017). The resulting plasmid (pNRB860) was co-transformed into DKB6885 and DKB6899 with an 80nt oligo consisting of the 40bp upstream of the *RAD17* initiating codon and 40bp of sequence starting downstream from the *RAD17* stop codon.

### Antibodies

Dilutions of primary antibodies are as follows: specific-pure, anti-goat Dmc1, DKB#192 (1:800); anti-rabbit RFA2, DKB#197, a gift from Dr. Shinohara (1:1000). Secondary antibodies were diluted as such: donkey anti-rabbit Alexa Fluor 594 (1:1000, Invitrogen by ThermoFisher Scientific, A11062); chicken anti-goat Alexa Fluor 488 (1:1000, Invitrogen by ThermoFisher Scientific, A11034).

### Imaging

Images were collected on a Zeiss Axiovision 4.6 wide-field fluorescence microscope at 100X magnification. The same imaging parameters were used for all samples from a given experiment. ImageJ and CellProfiler software were used for quantitative analysis of focus staining intensity.

### Western blots

Total protein lysates were prepared by TCA precipitation using a previously described method (Knop *et al*. 1999). Approximately 20-30μl of the sample was loaded into 10-12% SDS-PAGE gels, run at 100-130V for as long as needed to separate proteins of interest and transferred to PVDF (Immobilin-P, Millipore #IPVH00010) overnight at 40V at 4°C, with buffer stirring. The membrane was then blocked by soaking for 60min in 1X TBS + 0.1% Tween + 5% dried milk and then with buffer containing appropriate dilutions of primary antibodies including anti-cMyc DKB#269 (U of C Core) at 1:500; mouse anti-PGK1, DKB#203 (abcam ab113687) at 1:6000; and rabbit anti-Hop2, DKB#143 (Pacific Immunology) at 1:1000. Secondary antibodies used from Cytiva (Thermo Fisher): anti-mouse HRP-conjugated (#45000679, diluted 1:5000), anti-rabbit HRP-conjugated (#45000682, diluted 1:5000). Following washing, blots were incubated with Western Lightning (Perkin Elmer, NEL105001). Imaging was done on a LAS 4000, using GE Image Quant software.

For phospho-site analysis, samples were loaded on a 6% acrylamide gel and run at 40mA for 4hrs at room temperature and then transferred in 1X transfer buffer + 20% methanol (25mM Tris Base, 200mM Glycine) to PVDF (Immobilin-P, Millipore #IPVH00010) overnight at 40V at 4°C, stirring. Membranes were then processed for Western analysis as described above.

### Exo1 Purification

200ml of a meiotic culture at 4hrs was spun at 1200xg for 5min and washed with 20ml cold water + 1X PMSF (see chart below). The cell pellet (approximately 1g) was washed again with 20ml cold BH0.15 (25mM Hepes (pH 8), 2mM MgCl2, 0.1mM EDTA, 0.5mM EGTA-KOH,15% glycerol, 0.1% Nonidet P40,150mM KCL, 1mM DTT for washing) + 1X protease and phosphatase inhibitors (see chart below). Cells were again pelleted and resuspended in 1ml of cold BH0.15 + 1X inhibitors and dripped into liquid nitrogen to freeze at −80°C. The cells were then broken in a Spex 6875 grinding mill (with liquid nitrogen) for 1hr. 1X protease and phosphatase inhibitors were added and the powder was allowed to thaw quickly on ice and then transferred to a 15ml corex glass tube and centrifuged in a Sorvall SS-34 rotor at 12,000xg for 30min. After spinning, more 1X PMSF was added and the supernatant was transferred into a Beckman ultra-clear tube (13×51mm) and spun in a SW50.1 swinging bucket rotor in the ultra centrifuge at 40,000 rpm for 25 min. The supernatant was removed and more 1X protease and phosphatase inhibitors were added. 40mg of the cleared lysate was added to c-myc magnetic beads (100μl slurry washed in TBST, Thermo Scientific #PI88842) and Buffer H 0.15 (+ 1X all inhibitors) up to a volume of 1ml and this mixture was incubated at 4°C, rotating for 3hrs. 1X PMSF was added every 35 min. A magnet was used to remove unbound protein and then the beads were washed 6X in Buffer H 0.15 (+ 1X all inhibitors). The beads were resuspended in 60μl 5X non-reducing buffer (0.3M Tris•HCl, 5% SDS, 50% glycerol and proprietary pink tracking dye, Thermo Scientific #39001) and boiled for 5-10min. A magnet was used to separate beads from protein sample and 3μl of 1M DTT was added. The samples were run on 10% acrylamide SDS-PAGE gels and relevant bands were excised and submitted for analysis.

| Group | Inhibitor | Concentration | Solvent | Dilution | Company | Catalog # |
| --- | --- | --- | --- | --- | --- | --- |
| **A** | Chymostatin | 10mg/ml | DMSO | 1000 | Cayman Chemical | 15114 |
|  | Leupeptin | 10mg/ml | DMSO | 1000 | Cayman Chemical | 14026 |
|  | Antipain | 10mg/ml | DMSO | 1000 | Cayman Chemical | 26705 |
|  | Pepstatin A | 10mg/ml | DMSO | 1000 | Cayman Chemical | 9000469 |
|  | E-64 | 10mg/ml | DMSO | 1000 | Cayman Chemical | 10007963 |
|  | Aprotinin | 10mg/ml | DMSO | 1000 | Cayman Chemical | 14716 |
| **B** | Microcystin-LR | 100mM | Ethanol | 200 | Cayman Chemical | 10007188 |
| **C** | AEBSF-Pefablock | 2mM | Water | 200 | Cayman Chemical | 14321 |
|  | N-Ethylmaleimide (NEM) | 1mM | Water | 200 | Selleck Chemicals | S3692 |
| **D** | Sodium orthovanadate | 0.4mM | Water | 20 | Sigma | 450243 |
|  | Sodium Fluoride | 10mM | Water | 20 | Sigma | 201154 |
|  | Beta-glycerophosphate | 4mM | Water | 20 | Sigma | G9422 |
|  | Sodium Pyrophosphate | 2mM | Water | 20 | Sigma | 221368 |
| **E** | PMSF | 100mM | Isopropanol | 500 | Roche | 236608 |

### Mass spectrometry

In-gel digestion and analysis by LC-MS/MS was carried out at PTM BIO LLC (Shevchenko *et al*. 2006). The following text is a verbatim description of the methods that was provided by the company and is quoted here with the permission of Jun Cai.

#### “In-gel Digestion

For in-gel tryptic digestion, gel pieces were destained in 50 mM NH_4_HCO_3_ in 50% ACN (v/v) until clear. Gel pieces were dehydrated with 100 μl of 100% ACN for 5 min, the liquid removed, and the gel pieces rehydrated in 10 mM DTT and incubated at 37°C for 60 min. Gel pieces were again dehydrated in 100% ACN, liquid was removed and gel pieces were rehydrated with 55 mM IAA. Samples were incubated at room temperature, in the dark for 45 min. Gel pieces were washed with 50 mM NH4HCO3 and dehydrated with 100% ACN. Gel pieces were rehydrated with 10 ng/μl trypsin resuspended in 50 mM NH4HCO3 on ice for 1 h. Excess liquid was removed and gel pieces were digested with trypsin at 37 °C overnight. Peptides were extracted with 50% ACN/5% FA, followed by 100% ACN. Peptides were dried to completion and resuspended in 2% ACN/0.1%FA.

#### LC-MS/MS Analysis

The tryptic peptides were dissolved in solvent A (0.1% formic acid, 2% acetonitrile/water), directly loaded onto a home-made reversed-phase analytical column (25-cm length, 100 μm i.d.). Peptides were separated with a gradient from increase from 6% to35% solvent B (0.1% FA in 90% ACN) over 22 min and climbing to 80% in 4 min then holding at 80% for the last 4 min, all at a constant flow rate of 600 nL/min on an EASY-nLC 1000 HPLC system, The resulting peptides were analyzed by Q Exactive^TM^ Plus (Thermo Fisher Scientific).The peptides were subjected to NSI source followed by tandem mass spectrometry (MS/MS) in Q Exactive^TM^ Plus (Thermo) coupled online to the HPLC. Intact peptides were detected in the Orbitrap at a resolution of 70,000. Peptides were selected for MS/MS using NCE setting as 28; ion fragments were detected in the Orbitrap at a resolution of 17,500. A data-dependent procedure that alternated between one MS scan followed by 20 MS/MS scans was applied for the top 20 precursor ions above a threshold ion count of 5000 in the MS survey scan with 15.0s dynamic exclusion. The electrospray voltage applied was 2.2 kV. Automatic gain control (AGC) was used to prevent overfilling of the orbitrap; 5E4 ions were accumulated for generation of MS/MS spectra. For MS scans, the m/z scan range was 350 to 1800.

#### Data Processing

The resulting MS/MS data were processed using Proteome Discoverer 1.3 Tandem mass spectra were searched against Saccharomyces cerevisiae database. Trypsin/P was specified as cleavage enzyme allowing up to 2 missing cleavages. Mass error was set to 10 ppm for precursor ions and 0.02 Da for fragment ions. Carbamidomethyl on Cys were specified as fixed modification and oxidation on Met, Acetylation on protein N-terminal, phosphorylation on Thr/Tyr/Ser was specified as variable modifications. Peptide confidence was set at high, and peptide ion score was set > 20.”

## Supplemental Figure Legends

**Figure S1.**
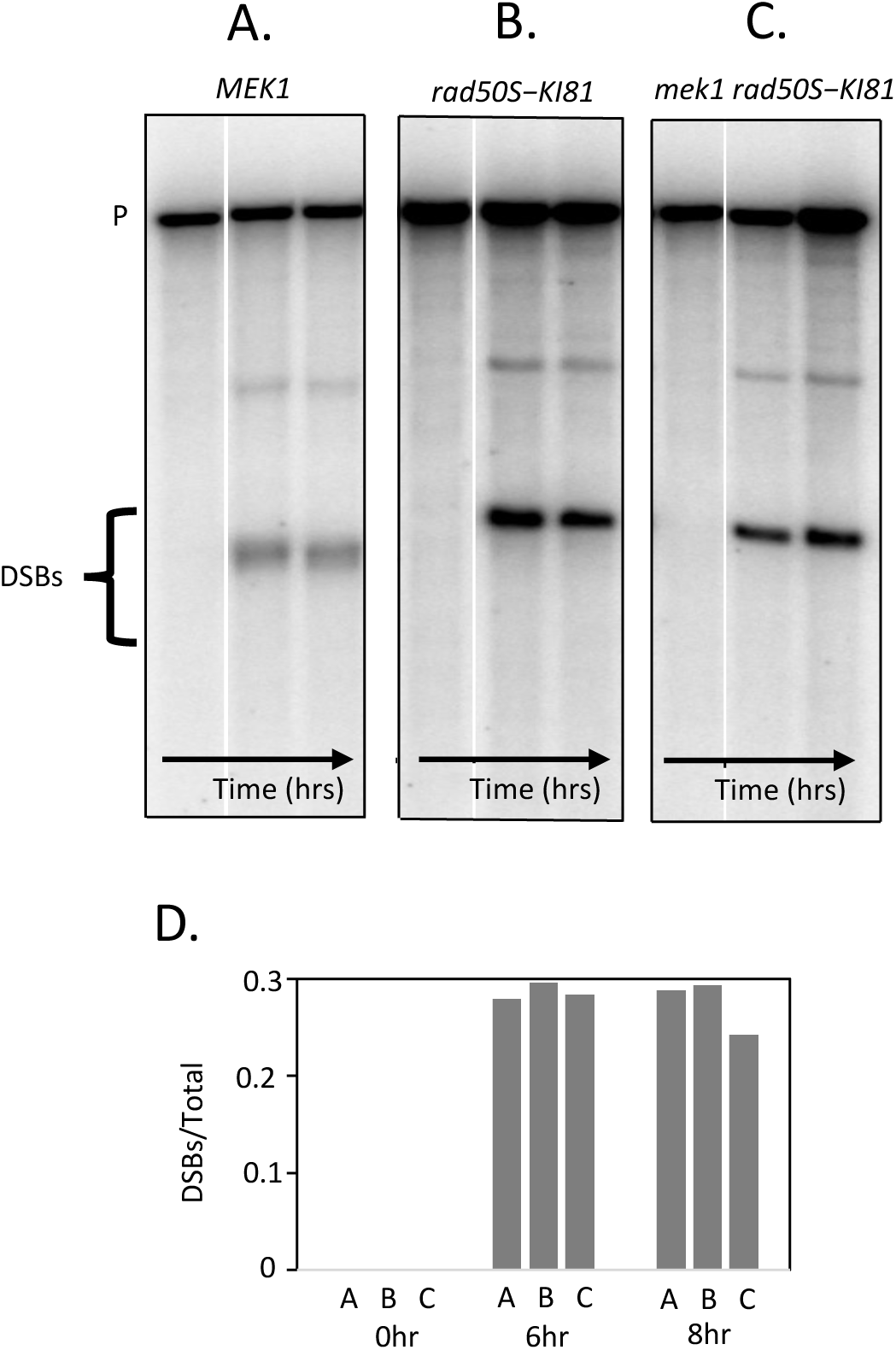
A *mek1Δ* mutation does not reduce DSB formation in *rad50S* cells. Southern analysis of DSB formation at the *HIS4::LEU2* hotspot. P=parental PstI fragment. DSBs=break fragments. Time points=0,6,and 8 hrs. **A.** *MEK1^+^, dmc1-II3A* (DKB6979) **B.** *MEK1^+^, dmc1-II3A, rad50-K181* (DKB6914) **C.** *mek1Δ, dmc1-II3A, rad50-K181* (DKB6913)

**Figure S2.**
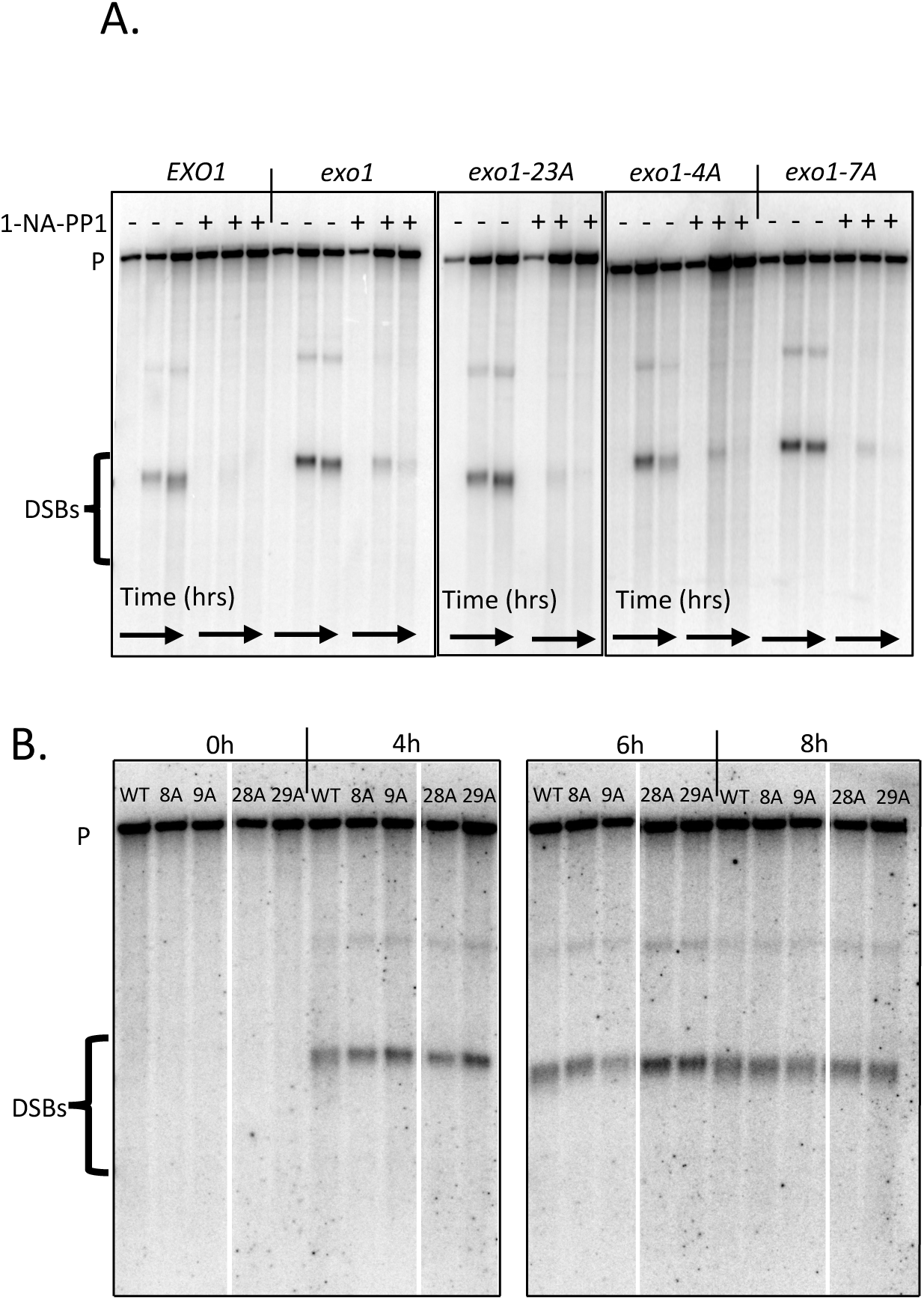
Additional analysis of Exo1 phosphosite mutants. Southern analysis of DSB formation at the *HIS4::LEU2* hotspot. P=parental PstI fragment. DSBs=break fragments. **A.** *EXO1-13XcMyc* (DKB6610), *exo1Δ* (DKB6620), *exo1-23A-13XcMyc* (DKB6764), *exo1-4S::A-13XcMyc* (DKB6611), *exo1-7A-13XcMyc* (DKB6613) Timepoints=0,4,6 hrs. 1µM 1-NA-PP1 or DMSO was added at time=0hrs. **B.** *EXO1-13XcMyc* (DKB6781), *exo1-8A-13XcMyc* (DKB7052), *exo1-9A-13XcMyc* (DKB7053), *exo1-28A-13XcMyc* (DKB7055), *exo1-29A-13XcMyc* (DKB7056). Time points=0,4,6 and 8hrs. No inhibitor was used.

**Table S1.**
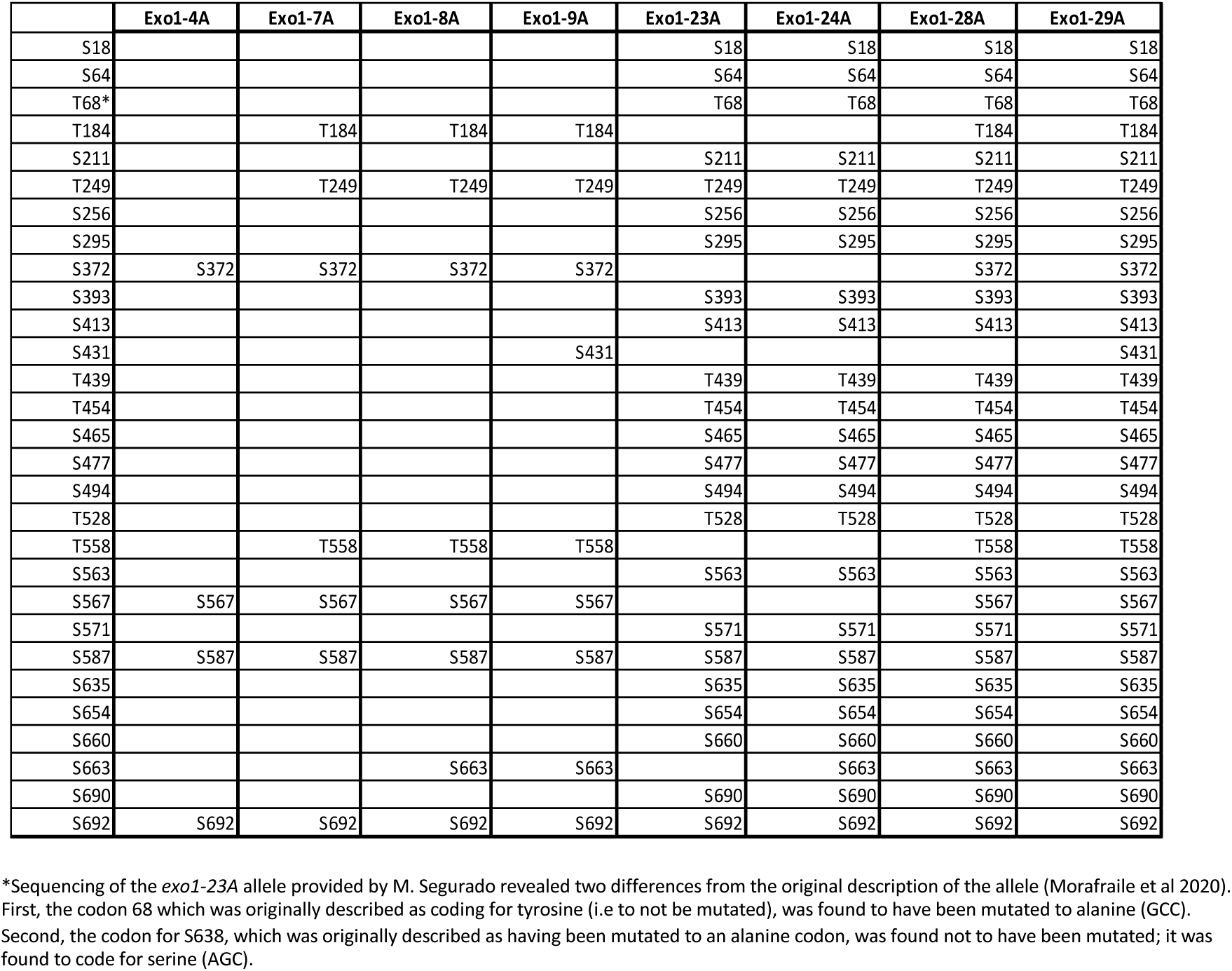

**Table S2.**
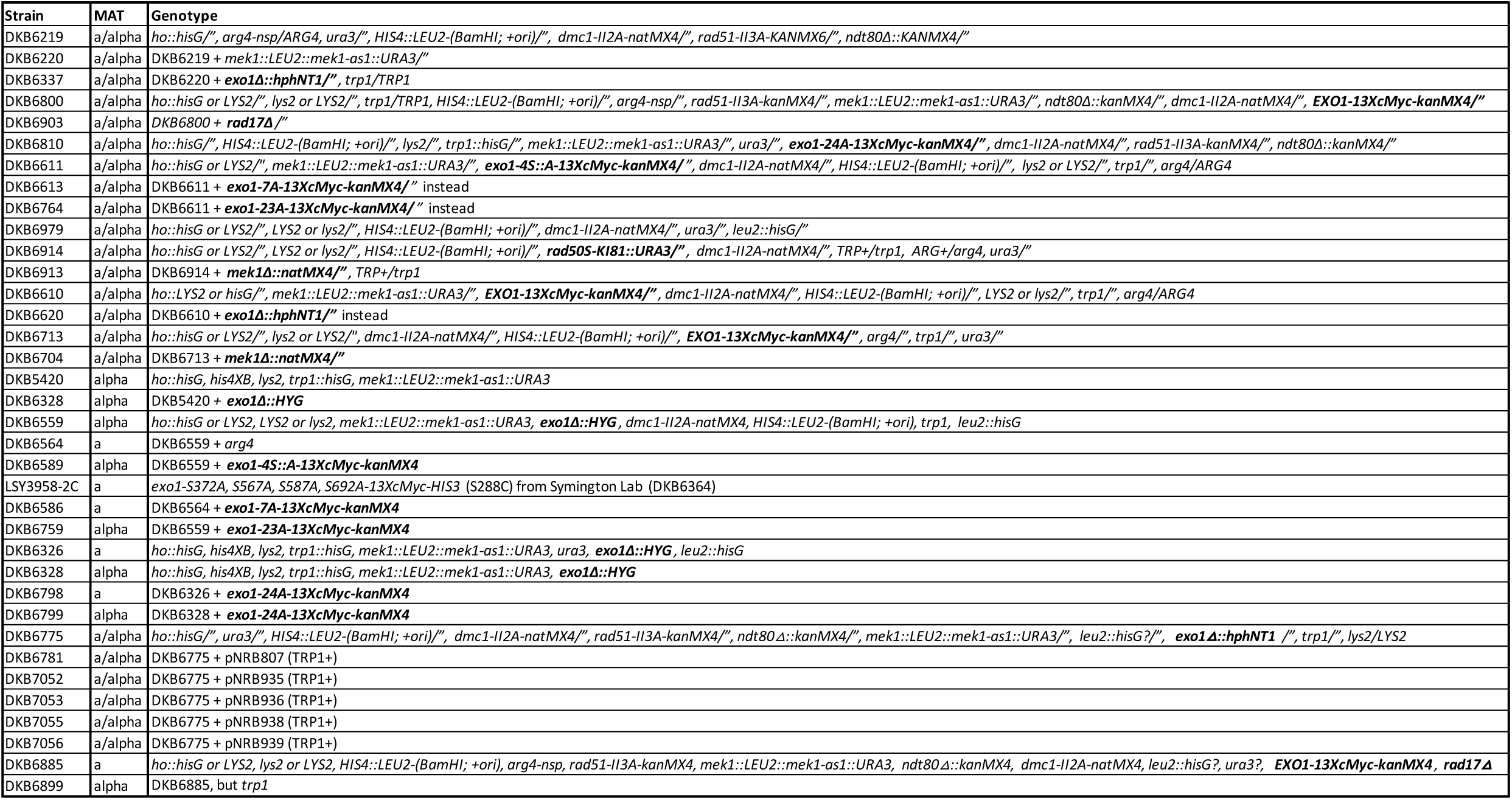

**Table S3.**

| plasmid name | Description |
| --- | --- |
| pRS41H | *hphNT1* (Taxis and Knop 2006) |
| pRS41K | *kanMX4* (Taxis and Knop 2006) |
| pNRB718 | *exo1* $\Delta$ *hphNT1* |
| pNRB721 | *exo1-4SA-13XcMyc-HIS3* |
| pNRB743 | *exo1-4SA-13XcMyc-kanMX4* |
| pNRB744 | *EXO1-13XcMyc-kanMX4* |
| pNRB749 | *exo1-7A-13XcMyc-kanMX4* |
| pNRB800 | *exo1-23A-13XcMyc-kanMX4* |
| pNRB803 | *exo1-24A-13XcMyc-kanMX4* |
| pNRB807 | *EXO1-13XcMyc-kanMX4, ARS4, CEN6, TRP1, AmpR* |
| pNRB811 | *exo1-24A-13XcMyc-kanMX4, ARS4, CEN6, TRP1, AmpR* |
| bRA89 | *PGK1-Cas9, hphNT1, ARS4, CEN6* (Anand et al. 2017) |
| pNRB860 | *PGK1-Cas9, RAD17-gRNA, hphNT1, ARS4, CEN6* |
| pNRB935 | *exo1-8A-13XcMyc-kanMX4, ARS4, CEN6, TRP1* |
| pNRB936 | *exo1-9A-13XcMyc-kanMX4, ARS4, CEN6, TRP1* |
| pNRB938 | *exo1-28A-13XcMyc-kanMX4, ARS4, CEN6, TRP1* |
| pNRB939 | *exo1-29A-13XcMyc-kanMX4, ARS4, CEN6, TRP1* |

## References

Bishop D. K., 1994 RecA homologs Dmc1 and Rad51 interact to form multiple nuclear complexes prior to meiotic chromosome synapsis. Cell 79: 1081–1092. 10.1016/0092-8674(94)90038-8

Bishop A. C., O. Buzko, and K. M. Shokat, 2001 Magic bullets for protein kinases. Trends Cell Biol 11: 167–172. 10.1016/s0962-8924(01)01928-6

Bolderson E., N. Tomimatsu, D. J. Richard, D. Boucher, R. Kumar, et al., 2010 Phosphorylation of Exo1 modulates homologous recombination repair of DNA double-strand breaks. Nucleic Acids Res 38: 1821–1831. 10.1093/nar/gkp1164

Callender T. L., R. Laureau, L. Wan, X. Chen, R. Sandhu, et al., 2016 Mek1 Down Regulates Rad51 Activity during Yeast Meiosis by Phosphorylation of Hed1. Plos Genet 12: e1006226. 10.1371/journal.pgen.1006226

Cao L., E. Alani, and N. Kleckner, 1990 A pathway for generation and processing of double-strand breaks during meiotic recombination in S. cerevisiae. Cell 61: 1089–1101. 10.1016/0092-8674(90)90072-m

Carballo J. A., A. L. Johnson, S. G. Sedgwick, and R. S. Cha, 2008 Phosphorylation of the Axial Element Protein Hop1 by Mec1/Tel1 Ensures Meiotic Interhomolog Recombination. Cell 132: 758–770. 10.1016/j.cell.2008.01.035

Chen X., R. T. Suhandynata, R. Sandhu, B. Rockmill, N. Mohibullah, et al., 2015 Phosphorylation of the Synaptonemal Complex Protein Zip1 Regulates the Crossover/Noncrossover Decision during Yeast Meiosis. Plos Biol 13: e1002329. 10.1371/journal.pbio.1002329

Chen X., R. Gaglione, T. Leong, L. Bednor, T. de los Santos, et al., 2018 Mek1 coordinates meiotic progression with DNA break repair by directly phosphorylating and inhibiting the yeast pachytene exit regulator Ndt80. Plos Genet 14: e1007832. 10.1371/journal.pgen.1007832

Cloud V., Y.-L. Chan, J. Grubb, B. Budke, and D. K. Bishop, 2012 Rad51 Is an Accessory Factor for Dmc1-Mediated Joint Molecule Formation During Meiosis. Science 337: 1222–1225. 10.1126/science.1219379

Fiorentini P., K. N. Huang, D. X. Tishkoff, R. D. Kolodner, and L. S. Symington, 1997 Exonuclease I of Saccharomyces cerevisiae Functions in Mitotic Recombination In Vivo and In Vitro. Mol Cell Biol 17: 2764–2773. 10.1128/mcb.17.5.2764

Gasior S. L., A. K. Wong, Y. Kora, A. Shinohara, and D. K. Bishop, 1998 Rad52 associates with RPA and functions with Rad55 and Rad57 to assemble meiotic recombination complexes. Genes Dev. 12: 2208–2221. 10.1101/gad.12.14.2208

Gioia M., L. Payero, S. Salim, G. F. V, A. F. Farnaz, et al., 2023 Exo1 protects DNA nicks from ligation to promote crossover formation during meiosis. Plos Biol 21: e3002085. 10.1371/journal.pbio.3002085

Gnügge R., and L. S. Symington, 2021 DNA end resection during homologous recombination. Curr Opin Genet Dev 71: 99–105. 10.1016/j.gde.2021.07.004

Grubb J., M. S. Brown, and D. K. Bishop, 2015 Surface Spreading and Immunostaining of Yeast Chromosomes. J Vis Exp Jove e53081. 10.3791/53081

Hollingsworth N. M., and R. Gaglione, 2019 The meiotic-specific Mek1 kinase in budding yeast regulates interhomolog recombination and coordinates meiotic progression with double-strand break repair. Curr Genet 65: 631–641. 10.1007/s00294-019-00937-3

Hunter N., 2011 Double duty for Exo1 during meiotic recombination. Cell Cycle 10: 2607–2609. 10.4161/cc.10.16.16452

Kim K. P., B. M. Weiner, L. Zhang, A. Jordan, J. Dekker, et al., 2010 Sister Cohesion and Structural Axis Components Mediate Homolog Bias of Meiotic Recombination. Cell 143: 924– 937. 10.1016/j.cell.2010.11.015

Knop M., K. Siegers, G. Pereira, W. Zachariae, B. Winsor, et al., 1999 Epitope tagging of yeast genes using a PCR-based strategy: more tags and improved practical routines. Yeast 15: 963– 972. 10.1002/(sici)1097-0061(199907)15:10b<;963::aid-yea399>3.0.co;2-w

Lao J. P., S. Tang, and N. Hunter, 2013 Native/Denaturing two-dimensional DNA electrophoresis and its application to the analysis of recombination intermediates. Methods Mol Biology Clifton N J 1054: 105–20. 10.1007/978-1-62703-565-1_6

Longtine M. S., A. M. III, D. J. Demarini, N. G. Shah, A. Wach, et al., 1998 Additional modules for versatile and economical PCR-based gene deletion and modification in Saccharomyces cerevisiae. Yeast 14: 953–961. 10.1002/(sici)1097-0061(199807)14:10<;953::aid-yea293>3.0.co;2-u

Lydall D., Y. Nikolsky, D. K. Bishop, and T. Weinert, 1996 A meiotic recombination checkpoint controlled by mitotic checkpoint genes. Nature 383: 840–843. 10.1038/383840a0

Manfrini N., I. Guerini, A. Citterio, G. Lucchini, and M. P. Longhese, 2010 Processing of Meiotic DNA Double Strand Breaks Requires Cyclin-dependent Kinase and Multiple Nucleases*. J. Biol. Chem. 285: 11628–11637. 10.1074/jbc.m110.104083

Mimitou E. P., S. Yamada, and S. Keeney, 2017 A global view of meiotic double-strand break end resection. Science 355: 40–45. 10.1126/science.aak9704

Morafraile E. C., A. Bugallo, R. Carreira, M. Fernández, C. Martín-Castellanos, et al., 2020a Exo1 phosphorylation inhibits exonuclease activity and prevents fork collapse in rad53 mutants independently of the 14-3-3 proteins. Nucleic Acids Res 48: 3053–3070. 10.1093/nar/gkaa054

Morafraile E. C., A. Bugallo, R. Carreira, M. Fernández, C. Martín-Castellanos, et al., 2020b Exo1 phosphorylation inhibits exonuclease activity and prevents fork collapse in rad53 mutants independently of the 14-3-3 proteins. Nucleic Acids Res 48: 3053–3070. 10.1093/nar/gkaa054

Morin I., H.-P. Ngo, A. Greenall, M. K. Zubko, N. Morrice, et al., 2008a Checkpoint-dependent phosphorylation of Exo1 modulates the DNA damage response. Embo J 27: 2400–2410. 10.1038/emboj.2008.171

Morin I., H.-P. Ngo, A. Greenall, M. K. Zubko, N. Morrice, et al., 2008b Checkpoint-dependent phosphorylation of Exo1 modulates the DNA damage response. Embo J 27: 2400–2410. 10.1038/emboj.2008.171

Niu H., L. Wan, B. Baumgartner, D. Schaefer, J. Loidl, et al., 2005 Partner Choice during Meiosis Is Regulated by Hop1-promoted Dimerization of Mek1. Mol Biol Cell 16: 5804–5818. 10.1091/mbc.e05-05-0465

Niu H., X. Li, E. Job, C. Park, D. Moazed, et al., 2007 Mek1 Kinase Is Regulated To Suppress Double-Strand Break Repair between Sister Chromatids during Budding Yeast Meiosis. Mol Cell Biol 27: 5456–5467. 10.1128/mcb.00416-07

Niu H., L. Wan, V. Busygina, Y. Kwon, J. A. Allen, et al., 2009 Regulation of Meiotic Recombination via Mek1-Mediated Rad54 Phosphorylation. Mol Cell 36: 393–404. 10.1016/j.molcel.2009.09.029

Oh S. D., J. P. Lao, A. F. Taylor, G. R. Smith, and N. Hunter, 2008 RecQ Helicase, Sgs1, and XPF Family Endonuclease, Mus81-Mms4, Resolve Aberrant Joint Molecules during Meiotic Recombination. Mol. Cell 31: 324–336. 10.1016/j.molcel.2008.07.006

Piazza A., W. D. Wright, and W.-D. Heyer, 2017 Multi-invasions Are Recombination Byproducts that Induce Chromosomal Rearrangements. Cell 170: 760–773.e15. 10.1016/j.cell.2017.06.052

Reitz D., J. Grubb, and D. K. Bishop, 2019 A mutant form of Dmc1 that bypasses the requirement for accessory protein Mei5-Sae3 reveals independent activities of Mei5-Sae3 and Rad51 in Dmc1 filament stability. Plos Genet 15: e1008217. 10.1371/journal.pgen.1008217

Roeder G. S., and J. M. Bailis, 2000 The pachytene checkpoint. Trends Genet. 16: 395–403. 10.1016/s0168-9525(00)02080-1

Sanchez A., C. Adam, F. Rauh, Y. Duroc, L. Ranjha, et al., 2020 Exo1 recruits Cdc5 polo kinase to MutLγ to ensure efficient meiotic crossover formation. Proc National Acad Sci 117: 30577– 30588. 10.1073/pnas.2013012117

Sertic S., R. Quadri, F. Lazzaro, and M. Muzi-Falconi, 2020 EXO1: A tightly regulated nuclease. Dna Repair 93: 102929. 10.1016/j.dnarep.2020.102929

Shevchenko A., H. Tomas, J. Havli, J. V. Olsen, and M. Mann, 2006 In-gel digestion for mass spectrometric characterization of proteins and proteomes. Nat. Protoc. 1: 2856–2860. 10.1038/nprot.2006.468

Shinohara A., H. Ogawa, and T. Ogawa, 1992 Rad51 protein involved in repair and recombination in S. cerevisiae is a RecA-like protein. Cell 69: 457–470. 10.1016/0092-8674(92)90447-k

Suhandynata R. T., L. Wan, H. Zhou, and N. M. Hollingsworth, 2016 Identification of Putative Mek1 Substrates during Meiosis in Saccharomyces cerevisiae Using Quantitative Phosphoproteomics. Plos One 11: e0155931. 10.1371/journal.pone.0155931

Symington L. S., 2016 Mechanism and regulation of DNA end resection in eukaryotes. Crit Rev Biochem Mol 51: 195–212. 10.3109/10409238.2016.1172552

Szankasi P., and G. R. Smith, 1992 A DNA exonuclease induced during meiosis of Schizosaccharomyces pombe. J Biol Chem 267: 3014–3023. 10.1016/s0021-9258(19)50688-3

Tomimatsu N., B. Mukherjee, M. C. Hardebeck, M. Ilcheva, C. V. Camacho, et al., 2014 Phosphorylation of EXO1 by CDKs 1 and 2 regulates DNA end resection and repair pathway choice. Nat Commun 5: 3561–3561. 10.1038/ncomms4561

Tsubouchi H., and G. S. Roeder, 2006 Budding yeast Hed1 down-regulates the mitotic recombination machinery when meiotic recombination is impaired. Gene Dev 20: 1766– 1775. 10.1101/gad.1422506

Wan L., T. de los Santos, C. Zhang, K. Shokat, and N. M. Hollingsworth, 2004 Mek1 Kinase Activity Functions Downstream of Red1 in the Regulation of Meiotic Double Strand Break Repair in Budding Yeast. Mol Biol Cell 15: 11–23. 10.1091/mbc.e03-07-0499

Xu L., M. Ajimura, R. Padmore, C. Klein, and N. Kleckner, 1995 NDT80, a Meiosis-Specific Gene Required for Exit from Pachytene in Saccharomyces cerevisiae. Mol. Cell. Biol. 15: 6572– 6581. 10.1128/mcb.15.12.6572

Xu L., B. M. Weiner, and N. Kleckner, 1997 Meiotic cells monitor the status of the interhomolog recombination complex. Genes Dev. 11: 106–118. 10.1101/gad.11.1.106

Yan S., S. Gao, and P. Zhou, 2021 Multi-functions of exonuclease 1 in DNA damage response and cancer susceptibility. Radiat Medicine Prot 2: 146–154. 10.1016/j.radmp.2021.08.004

Zakharyevich K., Y. Ma, S. Tang, P. Y.-H. Hwang, S. Boiteux, et al., 2010 Temporally and Biochemically Distinct Activities of Exo1 during Meiosis: Double-Strand Break Resection and Resolution of Double Holliday Junctions. Mol Cell 40: 1001–1015. 10.1016/j.molcel.2010.11.032

## Supplementary References

Anand R., G. Memisoglu, and J. Haber, 2017 Cas9-mediated gene editing in Saccharomyces cerevisiae. Protoc Exch. 10.1038/protex.2017.021a

Gietz R. D., and R. H. Schiestl, 2007 High-efficiency yeast transformation using the LiAc/SS carrier DNA/PEG method. Nat Protoc 2: 31–34. 10.1038/nprot.2007.13

Morafraile E. C., A. Bugallo, R. Carreira, M. Fernández, C. Martín-Castellanos, et al., 2020 Exo1 phosphorylation inhibits exonuclease activity and prevents fork collapse in rad53 mutants independently of the 14-3-3 proteins. Nucleic Acids Res 48: 3053–3070. 10.1093/nar/gkaa054

Taxis C., and M. Knop, 2006 System of centromeric, episomal, and integrative vectors based on drug resistance markers for Saccharomyces cerevisiae. BioTechniques 40: 73–78. 10.2144/000112040

